# Structural analysis of *de novo* designed binders targeting the closed state of HSP90

**DOI:** 10.64898/2026.06.03.729966

**Authors:** Dhiraj Srivastava, Sneha Singh, Kimberly Boyd, Nikolai O. Artemyev

## Abstract

Heat shock protein 90 (HSP90) assists protein folding and maturation of many important signaling proteins. In various diseases, HSP90 clients contribute to aberrant signaling, and HSP90 inhibition is being explored as a potential therapeutic approach. Commonly researched HSP90 inhibitors target the ATP-binding pocket, thereby disrupting the ATP-induced closure of HSP90. Drugs disrupting the HSP90 ATPase cycle by targeting the closed state of the chaperone have not been developed. Here, we present *de novo* design and selection of protein binders interacting exclusively with the closed state HSP90. Two such binders, H2 and H4, were identified that feature a similar fold and comparable affinities for HSP90 but display different binding kinetics. The structures of the HSP90 complexes with H2 and H4 were determined by cryo-EM single particle analysis, and they revealed high accuracy of the BindCraft predictions. H2 and H4 compete with p23 at one but not both symmetrical p23 binding sites on HSP90. H2 expressed in HEK293T cells moderately elevated expression of HSP70 and had no effect on the HSP90 level, suggesting muted heat shock response. Overall, this study demonstrates that the *de novo* binders represent novel and promising tools to probe the potential utility of HSP90 inhibition by targeting its closed state.

## INTRODUCTION

Heat shock protein 90 (HSP90) is a conserved molecular chaperone that plays a central role in regulating proteostasis by assisting protein folding and maturation, as well as affecting conformational dynamics of a diverse subset of proteins referred to as HSP90 clients [1-3]. In contrast to general chaperones that are involved in initial folding of nascent polypeptides, HSP90 preferentially interacts with and stabilizes or activates folded but metastable proteins. HSP90 clients include important signaling molecules such as kinases, transcription factors, E3 ligases, and hormone receptors [4-6]. HSP90 functions as an obligatory homodimer composed of three major domains: an N-terminal ATP-binding domain (NTD), a middle domain (MD) responsible for client interaction and catalytic regulation, and a C-terminal dimerization domain (CTD) [2]. Apo-HSP90 assumes an open extended or V-shape conformation in which the monomers are dimerized only through CTD. ATP binding to the NTD induces large-scale conformational changes, advancing N-terminal dimerization and formation of a closed state whereby a partially unfolded client protein is threaded through the lumen at the HSP90 dimer interface. Hydrolysis of ATP drives rearrangements in the dimer that induce activation/maturation of a client protein which is then released following dissociation of ADP and transition of HSP90 back to the open state [2]. At various steps of the ATPase cycle, HSP90 interacts with co-chaperones which modulate its ATPase activity and regulate client specificity. Beyond normal proteostasis, HSP90 is involved in pathogenesis of numerous diseases. HSP90 stabilizes many oncogenic client proteins responsible for proliferation, survival, and therapeutic resistance in cancer, including receptor tyrosine kinases (e.g. HER2, EGFR, MET), intracellular kinases (e.g. BRAF, RAF1, AKT, PI3K, CDK4), transcription factors (mutant p53, HIF-1, AR, ER) and aggregation-prone proteins such as tau and mutant huntingtin, underlying the chaperone’s involvement in neurodegenerative diseases [6-8].

Because of the broad stabilization of abnormal signaling pathways by HSP90, its pharmacological inhibition has become an attractive therapeutical strategy, which is being tested in clinical trials [6, 9]. Although efforts to target a second cryptic ATP-binding pocket in the HSP90 CTD as well as to target HSP90 interactions with co-chaperones have been described, by far the most studied and advanced HSP90 inhibitors target the N-terminal ATP-binding pocket, thereby disrupting the ATP-induced closure of HSP90 [6-9]. A common side-effect of the ATP-site inhibitors, which lock HSP90 in the open state, as potential therapeutics in cancer is the heat shock response (HSR) caused by release and activation of HSF1 [10]. HSF1-mediated upregulation of HSP40, HSP70 and HSP90, while potentially beneficial in neurodegenerative disorders, increases both survival and resistance to HSP90 inhibition of cancer cells [6]. Thus, development of novel inhibitors of HSP90 that evade HSR may be advantageous for the HSP90-inhibition-based therapies in cancer. Despite such potential, no inhibitors locking HSP90 in the closed state have been developed or characterized. Instead, the focus has been largely on targeting and inhibiting interactions of HSP90 with co-chaperones such as Cdc37, Aha1 and p23 [7-9, 11].

Recently, AI-based *de novo* design of protein binders has emerged as a powerful platform for precision therapeutics. Deep learning models such as AlphaFold, RoseTTAFold, and diffusion-based generative algorithms allow computational design of small and stable proteins that can bind to selected surfaces on target proteins [12-16]. Here, we explored the idea that conformation-specific protein binders for the closed state of HSP90 can be designed, with the validation of the accuracy of the prediction for the HSP90-binder complex by structural cryo-EM analysis.

## RESULTS

### Binders design

Binders against closed conformation of HSP90 were designed using Bindcraft [13]. Three hotspot residues (F15, I99, V167) from both chains of HSP90 dimer were selected to define the binding site using the structure of human HSP90β (PDB 8EOB) [17]. Total, 101 binders passing the default BindCraft validations and filtering criteria were designed. These binders were further validated by AlphaFold 3. For each binder-HSP90 complex, ten AlphaFold 3 models were generated and compared to the BindCraft design. All the binder folds were similar between the BindCraft and AlphaFold 3 models, but the AlphaFold 3 models for 6 binders showed binding sites and/or poses that were different from the original design. The remaining 95 binders were next examined for their ability to engage both monomers in the HSP90 dimer. The binding energies (dG_separated) for the interaction between binders and HSP90 were calculated using Rosetta InterfaceAnalyzer [18, 19]. Binders with binding energy for each chain of HSP90 between 35-65% of that for HSP90 dimer were selected (77 binders). Such selection was designed to reject binders appreciably interacting with the open state HSP90. The final selection of 31 binders for gene synthesis was made considering the BindCraft scoring metrics as well as diversification of the designs by excluding some of the similar binder folds (Suppl. Figs. 1,2) [13].

**Figure 1.**
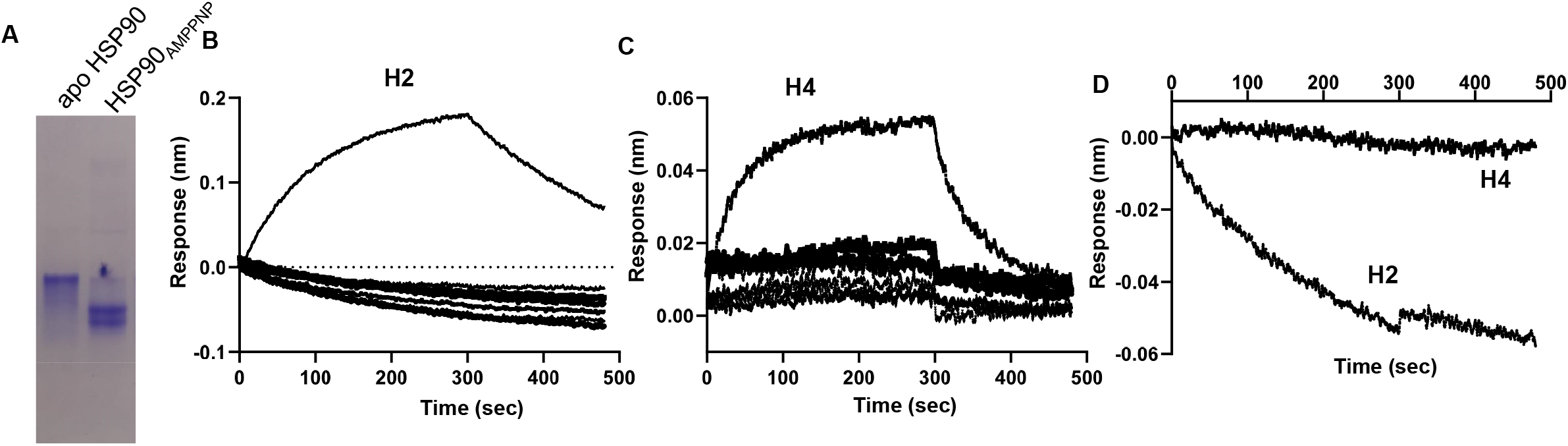
Screening of binders selective for the closed state HSP90. **A**. Coomassie-stained native gel to verify the closed state of HSP90 following binding of AMPPNP. The samples of apo HSP90 and HSP90_AMPPNP_ were prepared and subjected to native gel electrophoresis as described in *Methods*. HSP90_AMPPNP_ in the closed state conformation demonstrates higher mobility relative to the open state of apo HSP90. **B, C**. The Avi-tagged binders were immobilized to streptavidin coated biosensors using cleared lysates of autoinduced BL21(DE3) cells and the association and dissociation phase BLI responses were recorded for 7 binders in each run using 1 µM HSP90_AMPPNP_. Seven binders included H2 (B) and H4 (C). **D**. The BLI traces for H2 and H4 showing no binding to apo HSP90.

**Figure 2.**
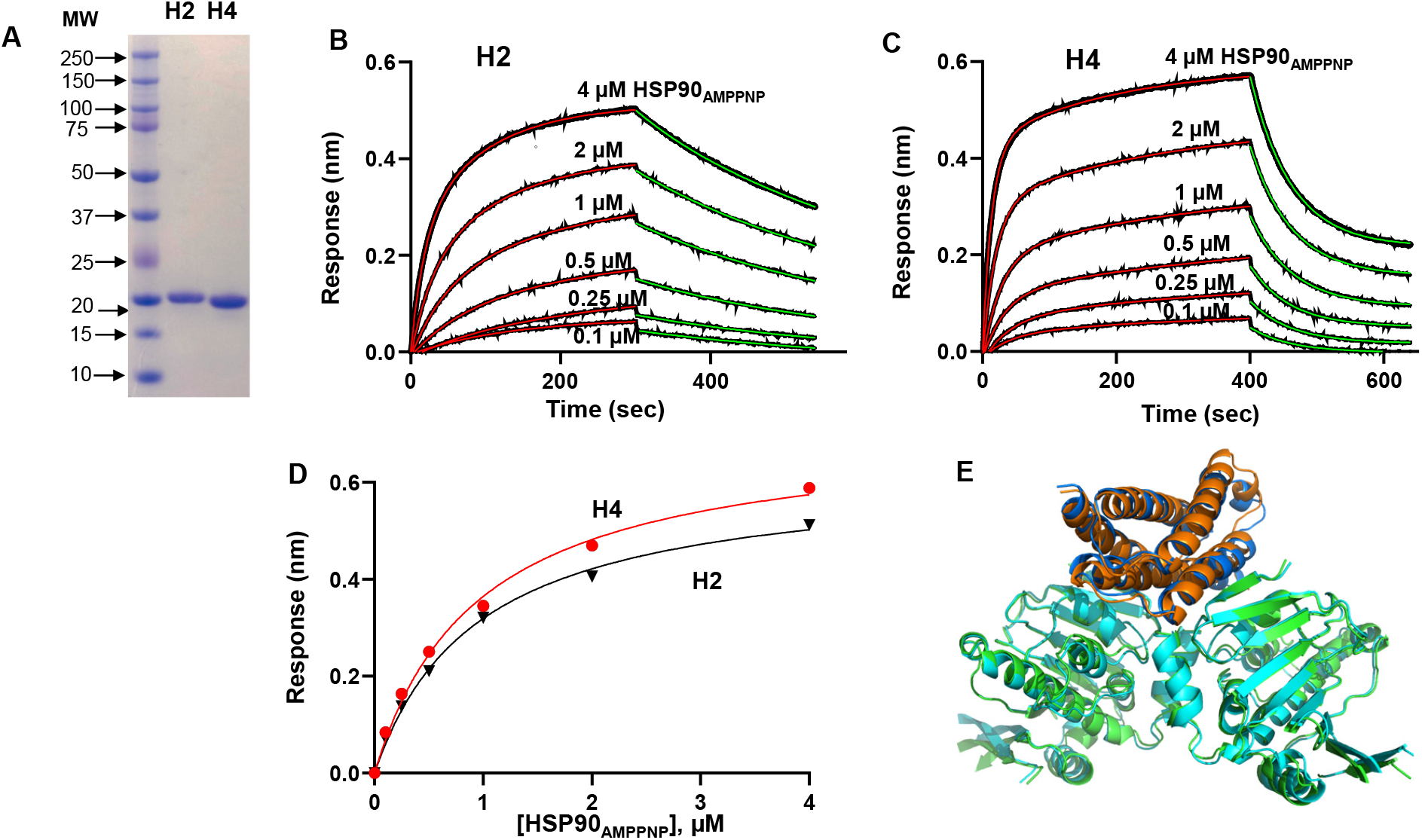
Interaction of HSP90 with H2 and H4. **A**. Coomassie-stained SDS-PAGE gel of purified preparations of H2 and H4. **B, C** Kinetics of association and dissociation for HSP90_AMPPNP_ and H2 (B) or H4 (C) coupled to a streptavidin biosensor as determined using BLI. The processed data curves are black and the nonlinear regression fits from the 1:1 binding model are red (association) and green (dissociation); **B**, k_on_=7.2±1.6 x 10^3^ M^-1^s^-1^; k_off_=2.4±0.3 x 10^-3^ s^-1^ (Mean±SD), K_D_(k_off_/k_on_)=0.33 µM; **C**, k_on_=1.9±0.2.6 x 10^4^ M^-1^s^-1^; k_off_=1.6±0.3 s^-2^, K_D_(k_off_/k_on_)=0.84 µM. (**D)** The steady state binding curves obtained from data in (B, C); H2, K_D_=0.94 µM; H4, K_D_=0.95 µM. For n=2 experiments, H2, K_D_=0.69 µM (range 0.44-0.94 µM); H4, K_D_= 0.93 µM (range 9.1-9.5 µM). **E**. Overlay of the BindCraft models for the HSP90 complexes with H2 (HSP90 - green, H2 – orange) and H4 (HSP90 - cyan, H2 -blue).

### Binder screening and characterization

Rapid screening of the computationally designed binders was conducted using bio-layer interferometry (BLI). The 31 selected binders were expressed each in a small volume bacterial culture and cleared cell lysates were used for BLI assays directly without additional purification. The Avi-tagged binders were immobilized to streptavidin coated biosensors. The association and dissociation phases were measured using 1 µM HSP90_AMPPNP_ or apo HSP90. The conversion of HSP90 from the open-state to the closed state during preincubation with AMPPNP was monitored using native gel PAGE (Fig. 1A). HSP90 has been shown to migrate faster in the closed-state compared to the open apo state during native gel PAGE [20]. Under our experimental conditions we routinely observed the appearance of two faster migrating bands apparently corresponding to the closed state HSP90. Although the exact nature of each of these bands is unclear, they may reflect known structural asymmetry of HSP90 protomers in the closed state [21]. The screening BLI experiments identified two genuine HSP90 binders, H2 and H4 (Fig. 1B,C). Binder H2 showed slower kinetics for both association and dissociation compared to H4 when assayed with the closed state HSP90_AMPPNP_ (Fig. 1B,C). Importantly, H2 and H4, did not bind the open-state apo HSP90, demonstrating selectivity of the interaction (Fig. 1D).

H2 and H4 were subsequently expressed on a larger scale and purified (Fig. 2A). Purified H2 and H4 were used for a quantitative characterization of their interaction with HSP90_AMPPNP_ by analyzing kinetics of association and dissociation at different concentrations of the chaperone and determining the binding affinity (K_D_) from the kinetic analysis as k_off_/k_on_ and the steady state data (Fig. 2). The mean K_D_ values for H2 and H4 from the kinetic analysis were 0.33 µM and 0.84 µM, respectively (Fig. 2 B,C) . The calculated K_D_ values for H2 and H4 from the steady-state analysis were somewhat higher, 0.69 µM and 0.93 µM, respectively (Fig. 2D). H2 and H4 originated from the same BindCraft initial step of binder backbone and sequence co-design, but the sequences diverged during the subsequent step of sequence optimization of non-interface regions [13]. Thus, the BindCraft models for the complexes of H2 and H4 with HSP90 feature similar folds (Fig. 2 E). **HSP90 binders H2 and H4 promote HSP90 transition to the closed state**.

We performed the native gel analysis of HSP90 in the absence or presence of H2 and H4 to examine if the binders alter migration of the closed state protein. Interestingly, the highly pure H2 binder (Fig. 2A), revealed several additional bands in native gel, possibly reflecting self-association (Fg. 3A). While H2 and H4 have a similar molecular size, H4 migrated significantly slower in the native gel compared to H2 (Fig. 3A,B). This observation is consistent with the difference in the pI values of for H2 and H4, respectively. Considering only ionizable side chains, H2 is 6-charge more negative than H4 at the native gel pH of 8.3 (Fig. 3C). As expected, addition of H2 or H4 to apo HSP90 did not affect the chaperone migration (Fig. 3A,B). Furthermore, H2 did not appreciably alter the mobility of the two bands corresponding to HSP90_AMPPNP_ (Fig. 3A). The lack of the noticeable effect of H2 is not surprising, given the mass change of ∼10% upon the complex formation with the binder. In contrast, the addition H4 to HSP90_AMPPNP_ shifted the two bands upwards (Fig. 3B). The effect of H4 may have been driven by its pI value. Notably, both H2 and H4 promoted the transition of HSP90 to the closed AMPPNP-bound state. This transition *in vitro* is not very efficient and typically requires incubation with ANPPNP for several hours [20], while leaving residual fraction of HSP90 in the open state. Incubation of HSP90 with AMPPNP in the presence of H2 or H4 led to a more efficient closure of HSP90 with no residual HSP90 in the open state (Fig. 3A,B).

**Figure 3.**
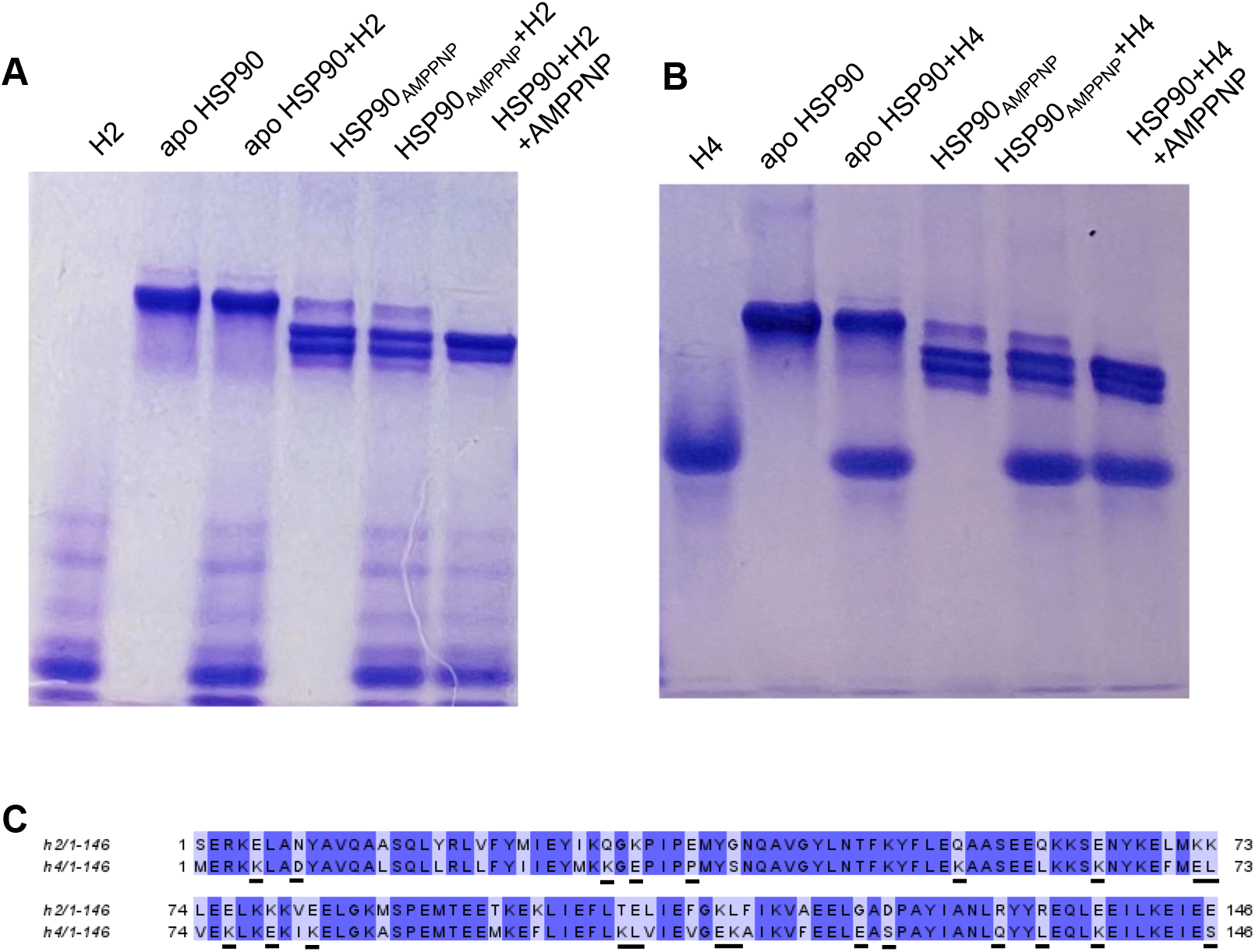
Native gel analysis of the interaction of HSP90 with the binders. **A, B**. Coomassie-stained native gels showing that apo HSP90 after incubation with AMPPNP adopts a faster migrating closed state. Addition of H2 did not alter the mobilities of apo HSP90 or HSP90_AMPPNP_ (**A**) whereas H4 had no effect on apo HSP90 but slowed the migration of both of the HSP90_AMPPNP_ bands (**B**). The efficiency of HSP90 closing increased when H2 or H4 were included into the AMPPNP incubation reaction (HSP90+H2+AMPPNP (**A**) and HSP90+H4+AMPPNP (**B**) samples). **C**. Sequence alignment for H2 and H4. Residues contributing to the charge difference of H2 and H4 are underlined.

Considering the difference in the overall charges of H2 and H4, we examined if it leads to different protein stabilities of the binders. The thermal stabilities of H2 and H4 were examined using dynamic light scattering (DLS). The DLS measurements demonstrated similar thermal stabilities of H2 and H4 with the onset of thermal aggregation at 51°C (Suppl. Fig. 3).

### Structures of the complexes of HSP90_AMPPNP_ with the H2 and H4 binders

For cryo-EM analysis, purified HSP90 was bound with AMPPNP in the presence of the binder proteins (H2 or H4) at 1:4 molar ratio and the samples were plunge frozen. The particles were picked using two different algorithms, Topaz picking and CryoSegNet [22, 23]. The two sets of particles for each of the H2 and H4 complexes were combined with removal of duplicates for further processing to increase the number of particles for final reconstructions. The consensus map of the HSP90**•**H2 complex based on 435.7 K particles was refined to an overall gold-standard FSC (GSFSC) resolution of 2.93 Å (Fig. 4A, Suppl. Figs. 4,6, Suppl. Table 1). The final reconstruction of the map for the HSP90**•**H4 complex based on 127.3 K particles was obtained at 3.06 Å GSFSC resolution (Fig. 4C. Suppl. Figs. 5,6, Suppl. Table 1). As indicated by the local resolution maps, the reconstructions featured well-resolved density for the NTD and MD of HSP90 and the portion of the binder involved in the interface with the chaperone, whereas the CTD and the binders’ residues opposite the interface were somewhat less resolved (Fig. 4B, D).

**Figure 4.**
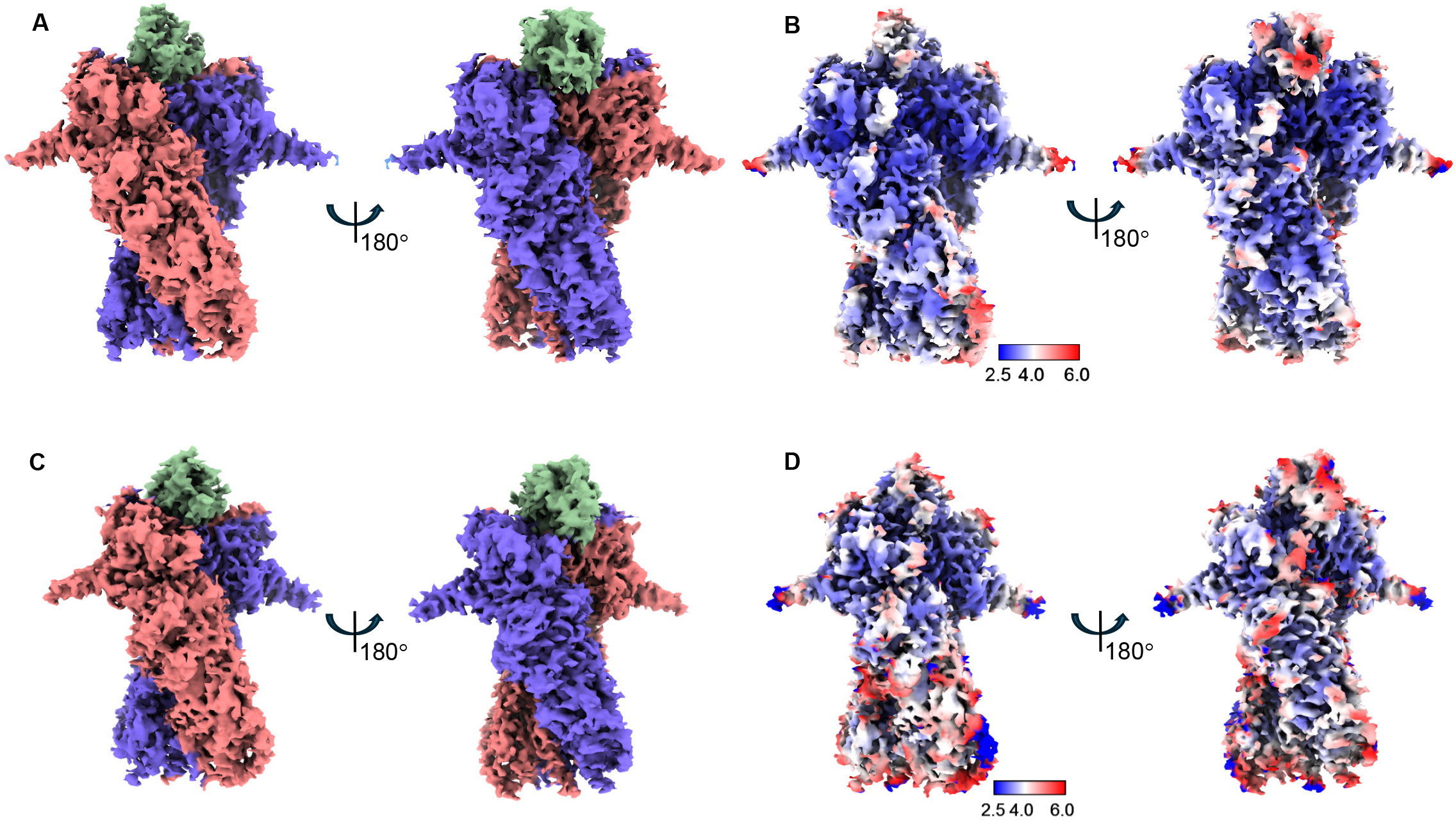
Cryo-EM maps for the complexes of HSP90 with H2 and H4 binders. The final cryoEM maps for the HSP90_AMPPNP_ complexes with H2 (A) and H4 (C) and the respective local resolution maps. The densities corresponding to the HSP90 subunits and the binders are colored in salmon, marine, and green.

**Figure 5.**
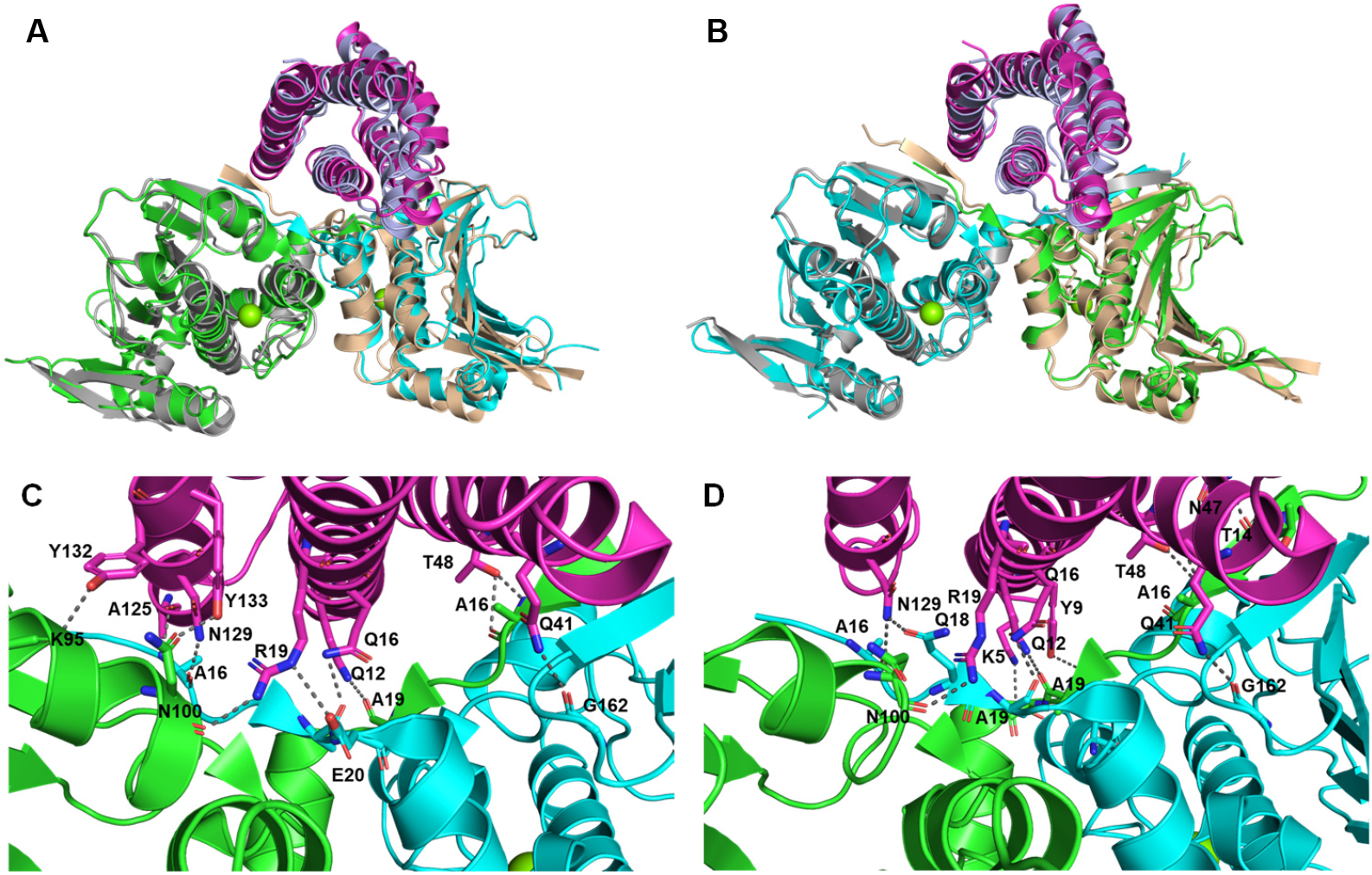
Structure models of the complexes of HSP90 with H2 and H4 binders. **A**,**B**. Overlay of the BinCraft binder designs H2 (**A**) and binder H4 (**B**) onto the respective cryoEM structures. The two chains of HSP90 are colored gray and wheat in the designs, and green and cyan in the cryo-EM structures. The binders are colored light blue in the designs, and magenta in the cryo-EM structure. Only the HSP90 NTDs with the binders are aligned and shown. The backbone RMSD between the designs and the cryo-EM structures are 1.78 Å (H2) and 1.22 Å (H4). **C, D**. Interactions between H2 (**C**) and H4 (**D**) and HSP90 based on the cryo-EM structures.

**Figure 6.**
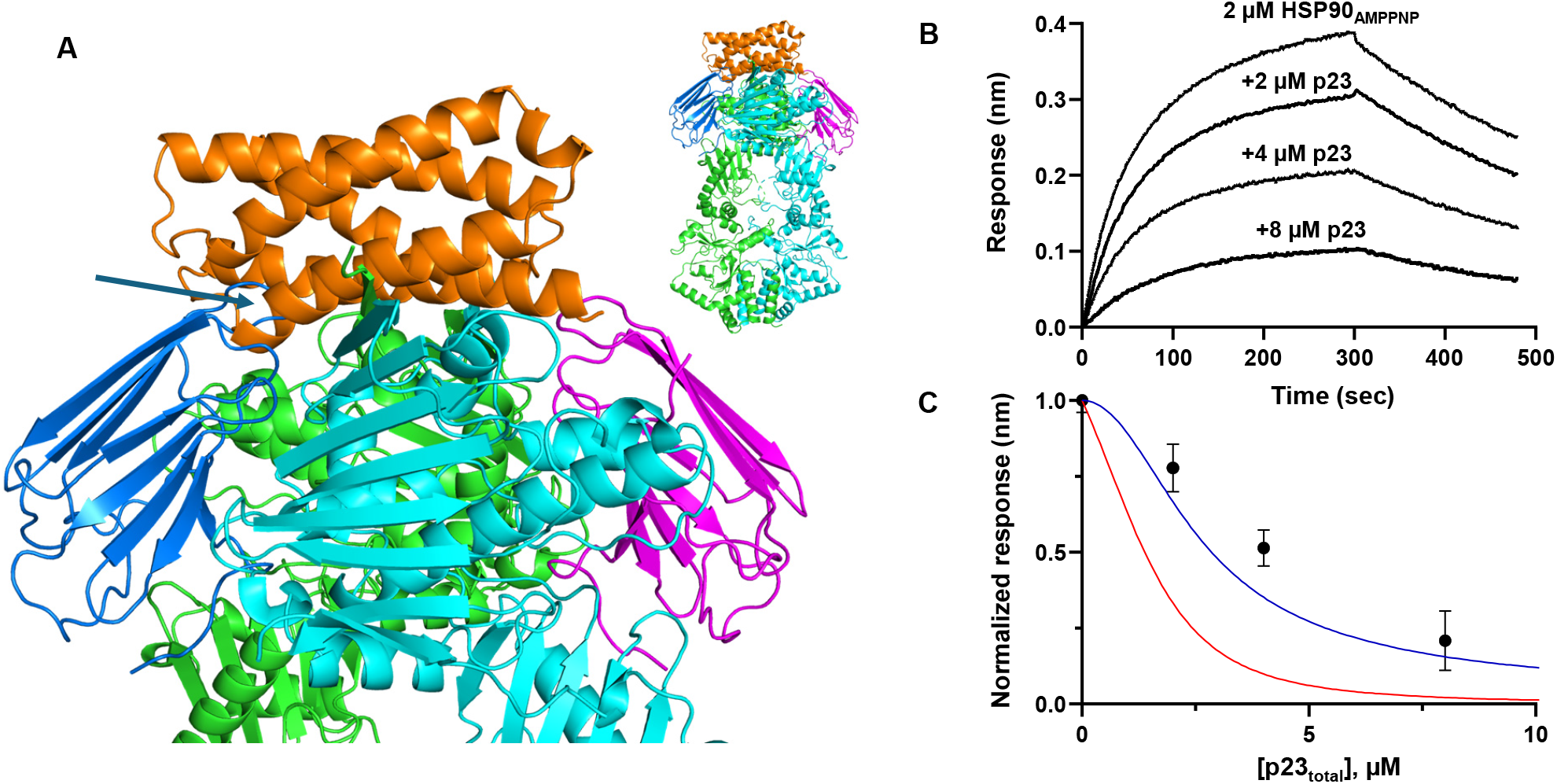
H2 allows P23 binding at one of the two symmetrical HSP90-binding sites. **A**. Two copies of the p23 molecules (PDB 7L7J) were superimposed on the structure of the HSP90/H2 complex. One p23 molecule (blue) shows a severe steric clash with H2, whereas the other molecule (magenta) can bind unobstructed. **B**. Representative BLI binding curves for H2 coupled to a streptavidin biosensor and HSP90_AMPPNP_ (2 µM) in the absence or presence of 2, 4, or 8 µM p23. **C**. Normalized BLI responses at 2, 4, and 8 µM of p23 as in (B) are shown as Mean±SE alone with the curves for the fractional concentrations of [HSP90]_free_ (red) and combined [HSP90]_free_ plus [HSP90**•**p23] (blue) simulated as described in *Methods*. The curves were simulated assuming the K_D_ value for the HSP90_AMPPNP_ **•**p23 complex of 1 µM [17].

Modeling the structures of the complexes of H2 and H4 with HSP90 into the final density maps indicated that BindCraft predicted the poses and specific interactions of H2 and H4 very accurately with the model to structure RMDS values of 1.78 Å and 1.22 Å (Fig. 5A,B). As predicted by BindCraft, H2 and H4 folded into an antiparallel 5-helical bundle. The structure of the H2/HSP90 complex shows that H2 acts as an interaction hub engaging the N-terminal domains of both chains A and B through a network of short-range contacts, including hydrogen bonds, electrostatic interactions, and hydrophobic packing interactions (Fig. 5C). The interface between H2 and chain A of HSP90 appears slightly stronger and more tightly packed than that with chain B. Notable H2/chain A contacts include hydrogen bonding interactions Y133(H2)–N100(A), T48(H2)–A16(A), and R19(H2)–N100(A). The interface is stabilized by additional polar contacts Y132(H2)–K95(A) and Y132(H2)–A96(A), and backbone-mediated interactions G44(H2)–T14(A) and T44(H2)–A16(A) contribute to packing. Hydrophobic packing is mediated by Y51(H2)-F15(A) side-chain stacking . H2 also engages chain B through several hydrogen-bonding and electrostatic interactions including N129(H2)–Q18(B), Q41(H2)–G162(B), R19(H2)–Q18(B) and R19(H2)–E20(B), and Q16(H2)–A19(B) and Q16(H2)–I21(B) (Fig. 5C). The interface of H4 with HSP90 is similar to that of H2. H4 lacks an ionic interaction between R19 and E20(B), but it has additional interactions compared to H2 (Fig. 5D). K5(H4) makes an ionic interaction with E20(A), whereas Y9(H4) sidechain makes a hydrogen bonding interaction with K107(B). Also, N47(H4) sidechain makes hydrogen bonding interactions with T14 (A) backbone carbonyl group (Fig. 5D).

### H2 allows P23 binding at one of the two symmetrical HSP90-binding sites

P23 is a well-characterized co-chaperone of HSP90 that selectively binds to the close-state conformation and inhibits ATP hydrolysis [20, 24-26]. There are two symmetrical p23 binding sites on HSP90 dimers although only one of these sites is apparently occupied in physiologically relevant complexes [2]. Overlay of the HSP90**•**H2 complex with the structure of the HSP90**•**p23 complex indicates that the bound H2 (or H4) would sterically interfere with one but not both p23 binding sites (Fig. 6A). We tested the interplay between p23 and H2 using the BLI binding assay where the ability of H2 to interact with HSP90_AMPPNP_ in the presence of p23 was examined (Fig. 6B). The assays were conducted with 2 µM HSP90_AMPPNP_. Under the BLI experimental conditions, the sensor bound H2 is a perfect “observer” of the binding species. Thus, the BLI responses in the presence of p23 normalized to that in the absence of p23 are expected to reflect fractional concentrations of the binding species. The normalized BLI responses in Fig. 6C suggest that both HSP90_free_ and its complex with a single copy of p23 bind H2 (Fig. 6C).

### H2 does not alter expression level of HSP90 transfected in HE293T cells

HSP90 inhibitors blocking its ATP-binding site induce the HSR leading to increased expression of HSPs including HSP70 and HSP90 [6, 7, 9, 27]. We tested if the designed binders modulate expression of HSP70 and HSP90 by transfecting HEK293T cells with the vector encoding H2. Transected cells were analyzed for expression of H2, HSP70, and HSP90 by Western blotting (Fig. 7). The immunoblot analysis demonstrated that H2 moderately elevated the expression level of HSP70. However, no changes in expression of HSP90 were detected (Fig. 7).

**Figure 7.**
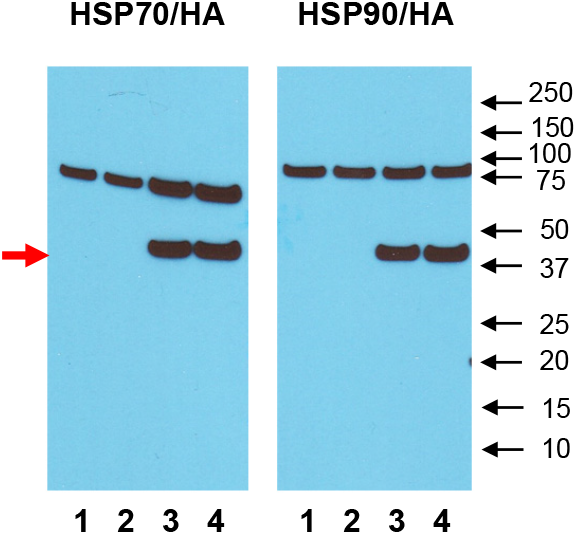
Effect of H2 on expression of HSP70 and HSP90 in HEK293T cells. HEK293T cells were mock transfected with the pcDNA3.1-Empty Vector (1 µg) (lane **1**), untransfected (lane **2**) or transfected with the pcDNA3.1 vector encoding the HA- and GFP-tagged H2 (1 µg and 2 µg; lanes **3** and **4**, respectively) and analyzed by Western blotting with anti-HSP70, HSP90, and HA antibodies. Red arrow indicates the band corresponding to HA-EGFP-H2.

## DISCUSSION

Inhibition of HSP90 with the drugs targeting its ATP-binding site is an active area of research with many clinical trials that have been completed or currently are underway [6-9, 11]. These drugs bind to HSP90 in the open state and block the ATPase cycle of the chaperone. Yet, such HSP90 inhibition through the accumulation of misfolded client proteins activates HSR, leading to increased expression of HSPs thereby counteracting the drug action in cancer [6]. Inhibitors of HSP90 selectively interacting with the closed-state HSP90 have not been described. This could be due to the difficulty of identifying druggable surfaces, as well as the uncertainty in the outcomes of stabilization of the closed state HSP90. Instead of blocking the HSP90 chaperone cycle, stabilization of the closed state could slow the cycle and enhance functional folding of oncogenes. Co-chaperone p23 preferentially binds and stabilizes closed HSP90 thereby extending client folding window and promoting maturation [20, 26, 28]. p23 is upregulated in multiple cancers and facilitates stabilization of oncogenic HSP90 clients [8, 11, 29, 30]. However, dead-end stabilization whereby HSP90 cannot return to the open state may potentially be beneficial due to misfolding of oncogenic clients, whereas HSR is muted because of accumulation of the closed state HSP90.

Assessing effects of stabilization of the closed state HSP90 for therapeutical applications requires novel experimental tools. We designed and validated two small protein binders specifically interacting with the closed-state HSP90. Two HSP90 binders, H2 and H4, originating from the common initial step of BindCraft backbone and sequence co-design were identified. The two binders feature a similar fold and the interaction interface with HSP90. Thus, similar binding affinities of H2 and H4 for HSP90_AMPPNP_ in the 0.3-0.9 µM range are not surprising. Nonetheless, the two binders revealed different association and dissociation kinetics of HSP90 binding suggesting a role for the binder surfaces outside the protein-protein interface. The cryo-EM structures of the HSP90 complexes with H2 and H4 indicated the high accuracy of the BindCraft prediction.

Analysis of the effect of H2 in cultured HEK293T cells indicated moderately increased expression of HSP70, but not HSP90 (Fig. 7). This observation suggests even though the binder’s inhibition of HSP90 does not escape the HSR, the latter might be muted in comparison to the response to the ATP-site HSP90 inhibitors. Additional utility of H2 and H4 is their potential to modulate the action of p23. Our analysis suggests that H2 and H4 compete with p23 only at one of the two symmetrical p23 binding sites of the closed state HSP90. This allows for the formation of a ternary complex HSP90/p23/H2(H4). Our previous analysis of p23 binding to HSP90AMPPNP indicated a very fast off-rate of ∼0.3 s^-1^ [17]. The dissociation rates for the binders, particularly for H2 (k_off_=2.4±0.3 x 10^-3^ s^-1^), are markedly lower indicating much longer lifetimes for the binder complexes with HSP90. Stabilization of the closed state HSP90 in the ternary complex is expected to increase the probability of the dead-end complex. Alternatively, given detrimental elevation of p23 in various tumors, the ability of binders to completely block p23 binding to HSP90 may become beneficial. Our structures suggest that modest modifications to the binders, such as a short extension of the N-terminal α-helix, would allow H2(H4) to block both p23 binding sites on HSP90. Overall, this study demonstrates the feasibility of *de-novo* design of site- and conformation specific binders of the closed-state HSP90. The ability to select the binding site and fine-tune binder interactions with HSP90 and co-chaperones should facilitate desirable outcomes in modulating the chaperone cycle. Novel small protein HSP90 binders may pave the way for design of new therapeutic strategies based on HSP90 inhibition.

## Methods

### HSP90 binders’ design

Binders against the closed conformation of HSP90 (PDB 8EOB) [17] were designed using the Bindcraft 4-stage design algorithm [13]. Three hotspot residues (F15, I99, V167) from both chains were used to define the binding site Total 101 binders were designed. Only residues 11-284 (excluding disordered residues 222-273) were used for binder design for reduce computational load. Binders passing the Bindcraft default validations and filtering criteria with initial guess-based complex prediction implemented for hard target were further validated using AlphaFold 3 [12]. Source code for AlphaFold 3 was obtained from google DeepMind GitHub repository and model parameters were obtained from Google DeepMind for non-commercial academic research. Input Json files for AlphaFold 3 were generated using a shell script and used full length HSP90 sequence, binders’ sequences, ATP and two random seeds generated using shell Random variable. Models of binder-HSP90 complexes were visually analyzed in PyMOL to ensure that AlphaFold 3 predictions were consistent with the Bindcraft design. Binders were further analyzed using Rosetta InterfaceAnalyzer and filtered based on binding energy (dG_separated) for the interaction between binders and HSP90 [18, 19]. Binders showing binding energy for each chain of HSP90 between 35-65 % of HSP90 dimer were selected. The final selection of binders was made based on the BindCraft scoring metrics such as pLDDT, iPTM and iPAE [13]. The genes were synthesized after conversion of protein sequences to nucleotide sequences and codon optimization for expression in *E. coli*. For expression in H293T cells, G block for H2, codon optimized for mammalian cell expression, was synthesized by IDT and cloned into a modified pCDNA 3.1 vector with N terminal HA and EGFP tag using Gibson assembly.

### Protein expression and purification

HSP90β was expressed and purified as described previously [17]. For the 31 selected binders, genes were synthesized in pET28a vector by TwistBioscience. The N-terminally Avi-tagged binders were transformed into BL21(DE3) cells and expressed in 2ml autoinduction media. The cells were grown at 37^°^C, harvested after 14-16 hours and lysed in 50mM TRIS, 200mM NaCl, 1mM TCEP, 5% glycerol, cOmplete Protease Inhibitor Cocktail (Roche). The cleared lysates were used in screening for binders interacting with HSP90 in the closed state. For subsequent characterization and cryo EM sample preparation, H2 and H4 were expressed in BL21(DE3) cells in 50ml autoinduction media at 20^°^C overnight. The two binders were purified using Ni-NTA resin (Genscript) followed by gel filtration on a HiLoad 26/60 Superdex 75 pg column (GE Healthcare) using buffer 10 mM HEPES (pH 7.5), 200 mM NaCl, 1 mM TCEP, and 5% glycerol. P23 was cloned [17], exressed in BL21(DE3) cells, and purified by Ni-NTA chromatography followed by gel filtration using HiLoad 26/60 Superdex 75 pg column equilibrated with buffer containing 10 mM HEPES (pH 7.5), 200 mM NaCl, 1 mM TCEP and 5% glycerol.

### Native gel electrophoresis

The closed state of HSP90_AMPPNP_ and the effect of binders were examined by native gel. For the screening with the binders, the purified HSP90β (100uM) was incubated with 8mM AMPPNP in a buffer with 10mM HEPES, 500mM KCl, 1mM TCEP, 5% glycerol, 10mM MgCl_2_. The reaction was carried out at 37^°^C for 3 hours. To analyze the effect of binders on the closing transition of HSP90, 8 µM of HSP90 was incubated with 60 µM (∼7X) of H2 and H4 binders and 8 mM AMPPNP in the buffer containing 10mM HEPES, 500 mM KCl, 1mM TCEP, 5% glycerol, 10 mM MgCl_2_ at 37^°^C for 3 hours. For native gel analysis, the samples were mixed with 2X Native sample buffer (62.5 mM Tris-HCl, pH 6.8, 40% glycerol, 0.01% bromophenol blue) and ran on a native gel (6% Tris glycine) for 2 hours at 100V.

### Bio-layer interferometry

The steady state along with the association and disassociation kinetics of H2 and H4 and HSP90 in the open and closed state were recorded by an Octet RED96 system and streptavidin (SA)-coated biosensors (FortéBio, Menlo Park, CA). The N-terminally Avi-tagged binders were immobilized to the SA biosensor at a concentration of 30 µg/ml for 80 seconds. The primary screening of binders to identify the actual binders was performed using 1 µM HSP90_AMPPNM_. H2 and H4 binders were further tested for binding with the open state apo HSP90. Binding studies were performed at 26^°^C with the buffer containing 10 mM HEPES, 200 mM NaCl, 1 mM TCEP, 5% glycerol, 0.5 mg/ml BSA. The biosensors were stirred in 200 µl of sample in each well at 1000 rpm and the data was acquired at the rate of 5 Hz. For correction of the non-specific binding, reference sensors were used without the addition of binders. The average k_on_ and k_off_ were calculated as means of the individual k_on_ and k_off_ values for all curves. The affinity constant K_D_ was calculated as mean k_off_/ mean k_on_ or from fitting steady-state data to the equation for one site-specific binding using the GraphPad Prism 10 software. For the BLI binding reactions in the presence of p23, total concentration of HSP90_AMPPNM_ (A_T_) was 2 µM, whereas total concentration of p23 varied at 2, 4, and 8 µM. Decrease in fractional BLI response due to competition of p23 and H2 was determined. For the HSP90/p23 binding rection A+B ⇌ AB and AB+B ⇌ AB_2_, where both steps share the same dissociation constant K_D_, the fractional concentrations (Y) of [HSP90]_free_ and combined [HSP90]_free_ plus [HSP90**•**p23] are given by equations 1 and 2, respectively:

1. Y = 1 / (1 + (B_free_/K_D_) + (B_free_/K_D_)^2)
2. Y = (1 + B_free_/K_D_) / (1 + (B_free_/K_D_) + (B_free_/K_D_)^2);

where the free concentration of p32, B_free,_ is a function of (X) total concentration of p23: B_free_ =(X - A_T_-K_D_ + sqrt((X - A_T_-K_D_)^2 + 4*X*K_D_)) / 2. Equations for [HSP90]_free_ and combined [HSP90]_free_ plus [HSP90**•**p23] were simulated for A_T_=2 µM and K_D_ of 1 µM for the HSP90**•**p23 complex using the GraphPad Prism 10 software.

### Dynamic Light Scattering

Dynamic light scattering (DLS) was used to examine thermostability of HSP90 binders H2 and H4. Purified binders were tested at concentration 1 mg/ml. DLS data for thermostability measurements were collected while heating the samples from 20 °C to 80 °C at 1 °C/min. Onset of protein thermal unfolding (T_onset_) was determined by the sudden increase in hydrodynamic radius during the temperature ramp. Analyses of thermostability were performed using a DynaPro Nanostar instrument (Wyatt) and data were analyzed using the Dynamics 7.1.7 software.

### Cell culture, transfection and Western blotting

HEK293T cells were cultured and maintained in DMEM containing 10% FBS (Gibco). Cells were transfected with pcDNA3.1-Empty Vector (1 µg) or pcDNA3.1 vector encoding the HA- and GFP-tagged H2 in concentrations of 1 µg and 2 µg using FuGene 6 (Promega) according to the manufacturer’s instructions. Transfected cells were collected 70 hours after transfection. Cell lysates were prepared in 20 mM Tris-HCl buffer (pH 7.5) containing 120 mM KCl, 1 mM MgCl2, and 1× EDTA-free protease inhibitor cocktail (Roche), and they were analyzed by Western blotting for protein expression. Proteins were separated by 4%–12% SDS-PAGE and then transferred to a nitrocellulose membrane using the iBlot Western blot kit (Invitrogen). The blots were analyzed using mouse monoclonal anti-HA (BioLegend, clone 16B12, catalog 901502, 1:1000 dilution), rabbit polyclonal anti-HSP70 (Proteintech, 10995-1-AP, 1:5000 dilution), and rabbit polyclonal anti-HSP90 (Santa Cruz, SC-7947, 1:2000 dilution) primary antibodies. The antibody-antigen complexes were detected using anti-mouse IgG horseradish peroxidase linked antibody (Cell Signaling Technology, 7076S, 1:10,000 dilution), horseradish peroxidase–conjugated goat anti-rabbit secondary antibody (Santa Cruz, SC-2004, 1:10,000 dilution), and enhanced chemiluminescence reagents (GE Healthcare).

### Cryo-EM sample preparation, data collection, and processing

HSP90 was mixed with H2 and H4 binders in a molar ratio of 1:5 along with 8mM AMPPNP and 10 mM HEPES, 500 mM KCl, 1 mM TCEP, 5% glycerol, 10 mM MgCl_2_ at 37^°^C for 3 hours. The sample was purified using BioSpin Micro spin 6 column equilibrated with 10 mM HEPES, 100 mM KCl, 1 mM TCEP, 5 mM MgCl_2_. 3 µl of sample with a final concentration of 0.5 mg/ml was applied to the glow discharged quantifoil holey carbon grids (R1.2/1.3). The sample was blotted for 3 sec at 4^°^C and 100% humidity. The grids were plunge frozen in liquid ethane using FEI Vitrobot (Thermo Fisher). Samples were imaged on a Titan Krios TEM (ThermoFisher Scientific) equipped with a Gatan K3 direct electron detector at a magnification of 105,000 and physical pixel sizes of 0.825 Å (0.4125 Å super resolution mode). The movies were collected with 40 frames each with a defocus range of -1.0 to -3.0 μm and a total exposure of 50 e-/Å^2^. 11,234 and 4,366 movies were collected for the complexes with H2 and H4, respectively.

The cryo-EM data processing for all reconstructions was performed using cryoSPARC V4.7.1 [31]. The movies were motion-corrected using patch motion correction with binning by 2 followed by patch CTF estimation. Particle picking was conducted using Topaz and CryoSegNet (). A Topaz model was generated based on 7.2 K particles selected after several rounds of 2D classification of particles picked by Blob Picker from 100 micrographs from the HSP90**•**H2 dataset. The Topaz and CryoSegNet sets of particles for each of the H2 and H4 complexes were combined and the duplicates removed prior to further processing. Flow charts for cryo-EM data processing for the HSP90**•**H2 and HSP90**•**H4 complexes are shown in Suppl. Fig. 4 and Suppl. Fig. 5, respectively. For the HSP90**•**H4 data processing, 3D classification without alignment using a focused mask for the HSP90 NTDs and H4 was added to improve the binder density in the reconstruction.

## Acknowledgments

This work was supported by the National Institutes of Health grant RO1 EY-10843 to N.O.A. We would like to acknowledge use of resources at the Carver College of Medicine’s Protein and Crystallography Facility at the University of Iowa. A portion of this research was supported by NIH grant U24GM129547 and performed at the PNCC at OHSU and accessed through EMSL (grid.436923.9), a DOE Office of Science User Facility sponsored by the Office of Biological and Environmental Research.

## Author contributions

D.S., S.S, and K.B. performed the experiments; D.S., S.S and N.O.A. analyzed the data; N.O.A. designed and supervised the project, and wrote the manuscript with input from D.S. and S.S.

---

**Suppl. Figure 1.**
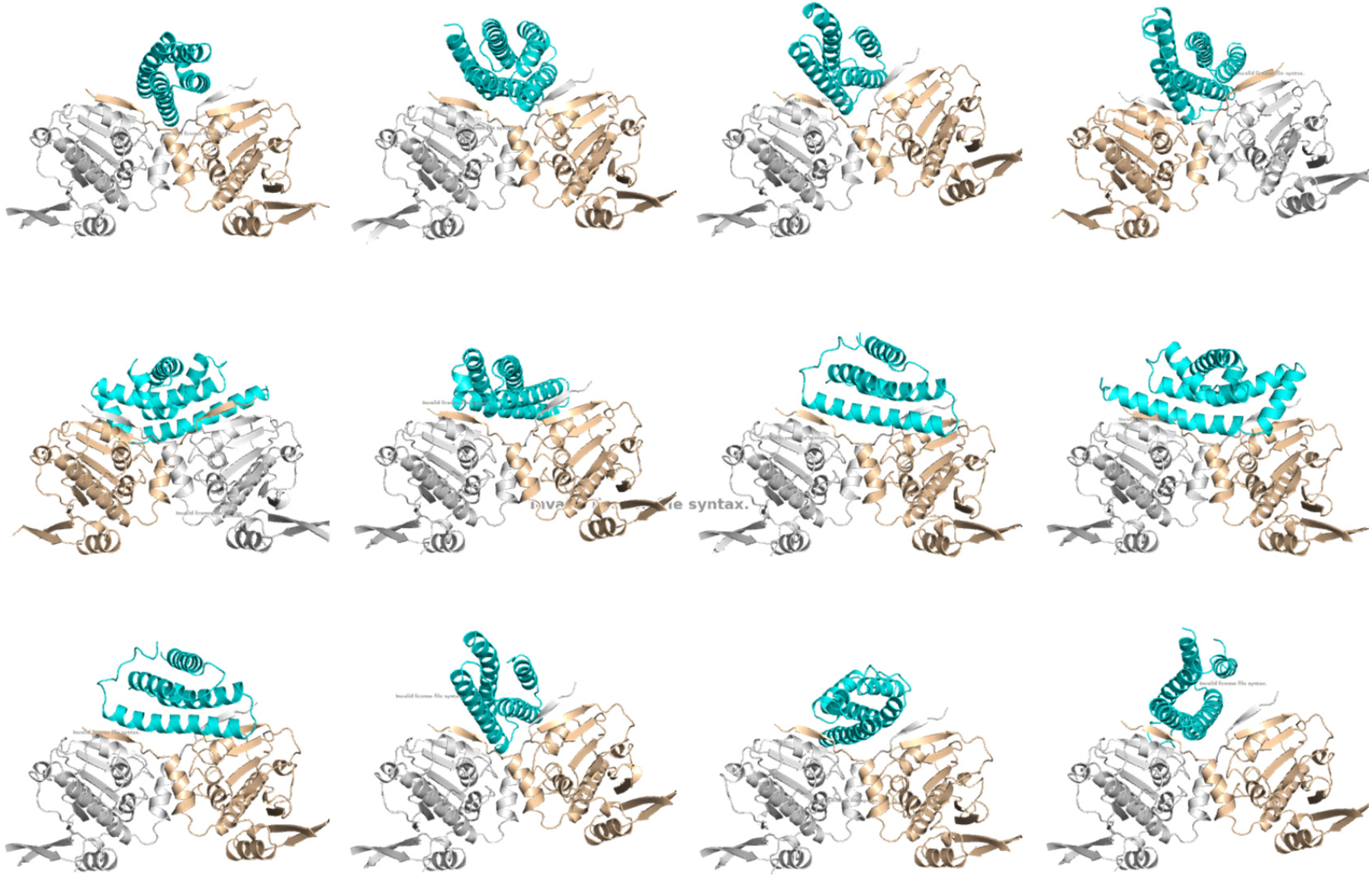
Top 12 BindCraft HSP90 binder designs.

**Suppl. Figure 2.**
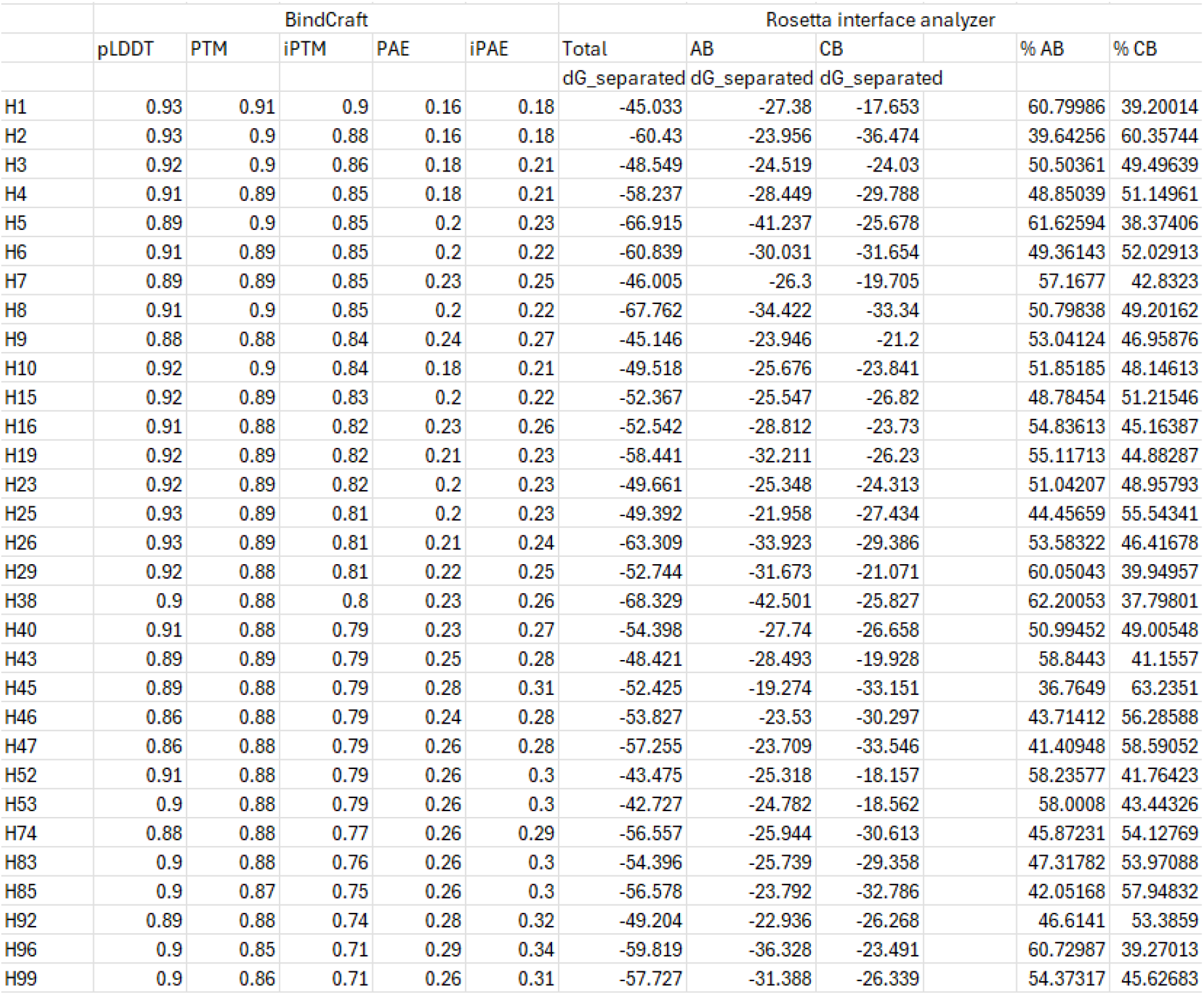
BindCraft and Rosetta Interface analyzer metrics for the selected 31 HSP90 binders.

**Suppl. Figure 3.**
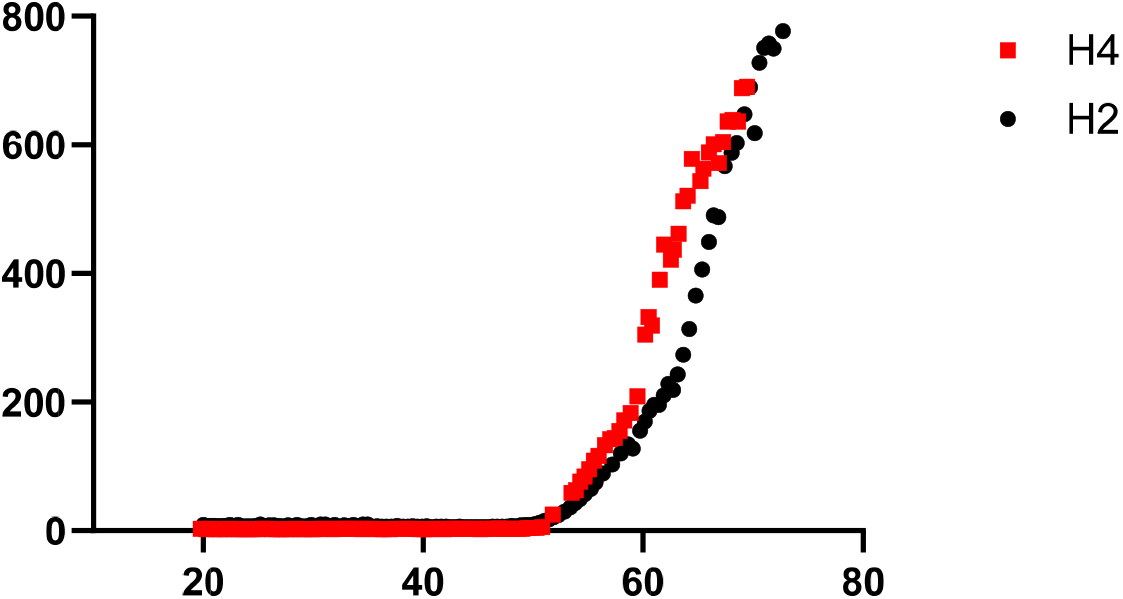
Thermal stabilities of H2 and H4. Representative thermal denaturation curves of H2 and H4 as determined by an increase in hydrodynamic radius on protein unfolding during a 1°C/min temperature ramp by dynamic light scattering. From three similar experiments, the Tonset values are (°C): H2, 51.3±0.4°C; H4, 51.0±0.2°C;

**Suppl. Figure 4.**
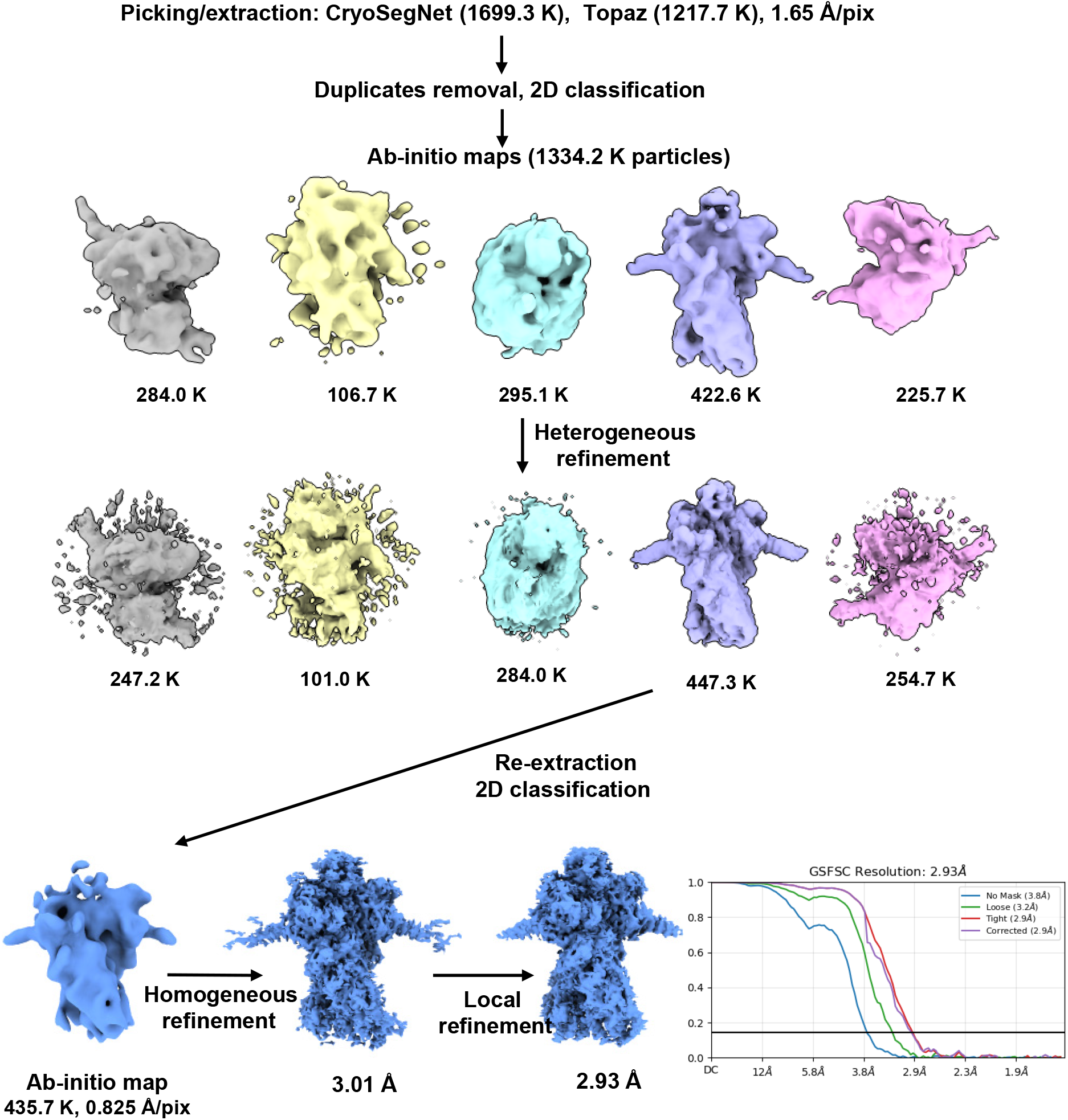
Flow chart of the cryo-EM analysis of the HSP90_AMPPNP_•H2 complex. Gold-Standard Fourier shell correlation (GSFSC) resolutions are shown for the final refinement steps. Fourier shell correlation (FSC) curves of the final map.

**Suppl. Figure 5.**
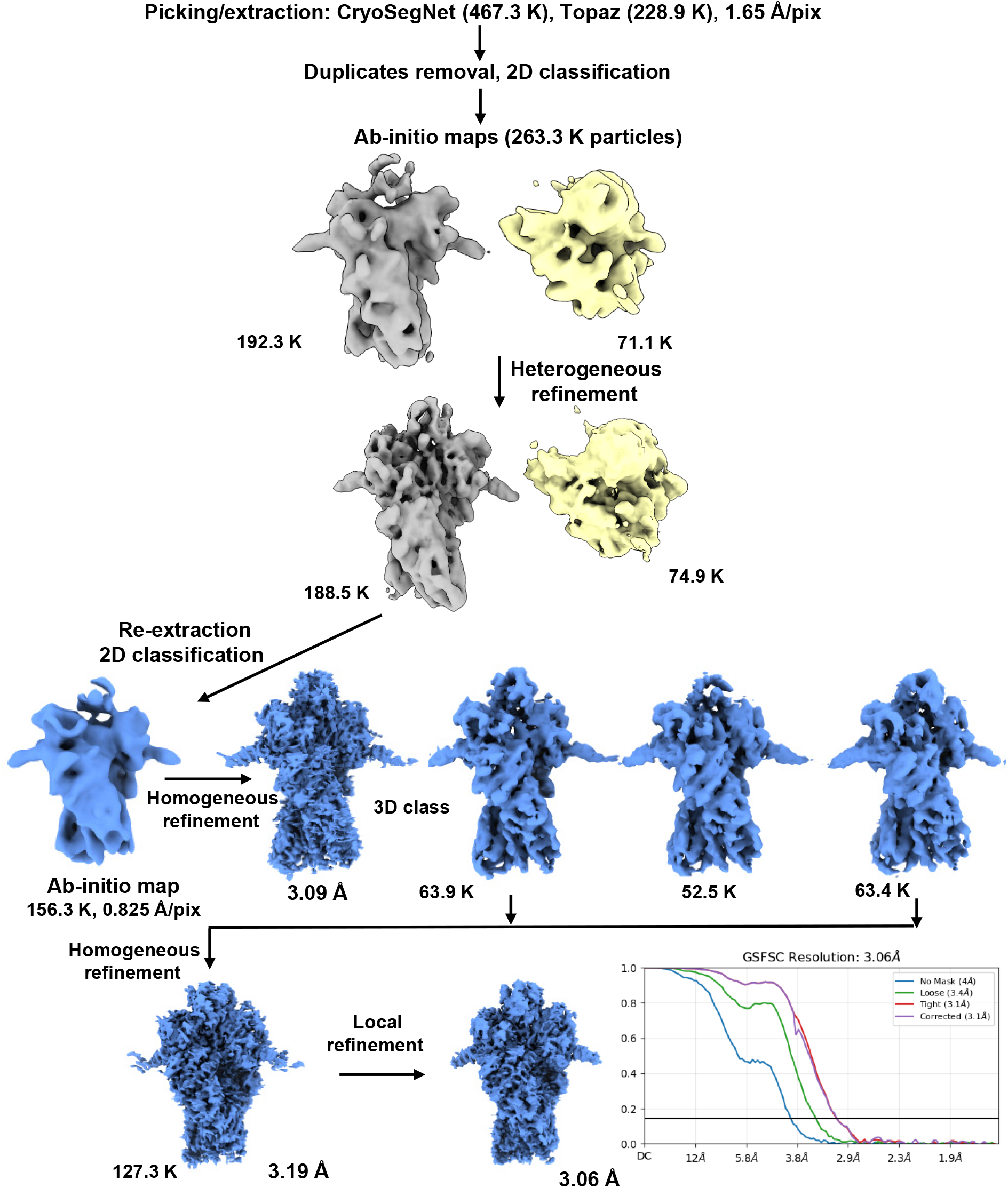
Flow chart of the cryo-EM analysis of the HSP90_AMPPNP_•H4 complex. Gold-Standard Fourier shell correlation (GSFSC) resolutions are shown for the late refinement steps. Fourier shell correlation (FSC) curves of the final map.

**Suppl. Figure 6.**
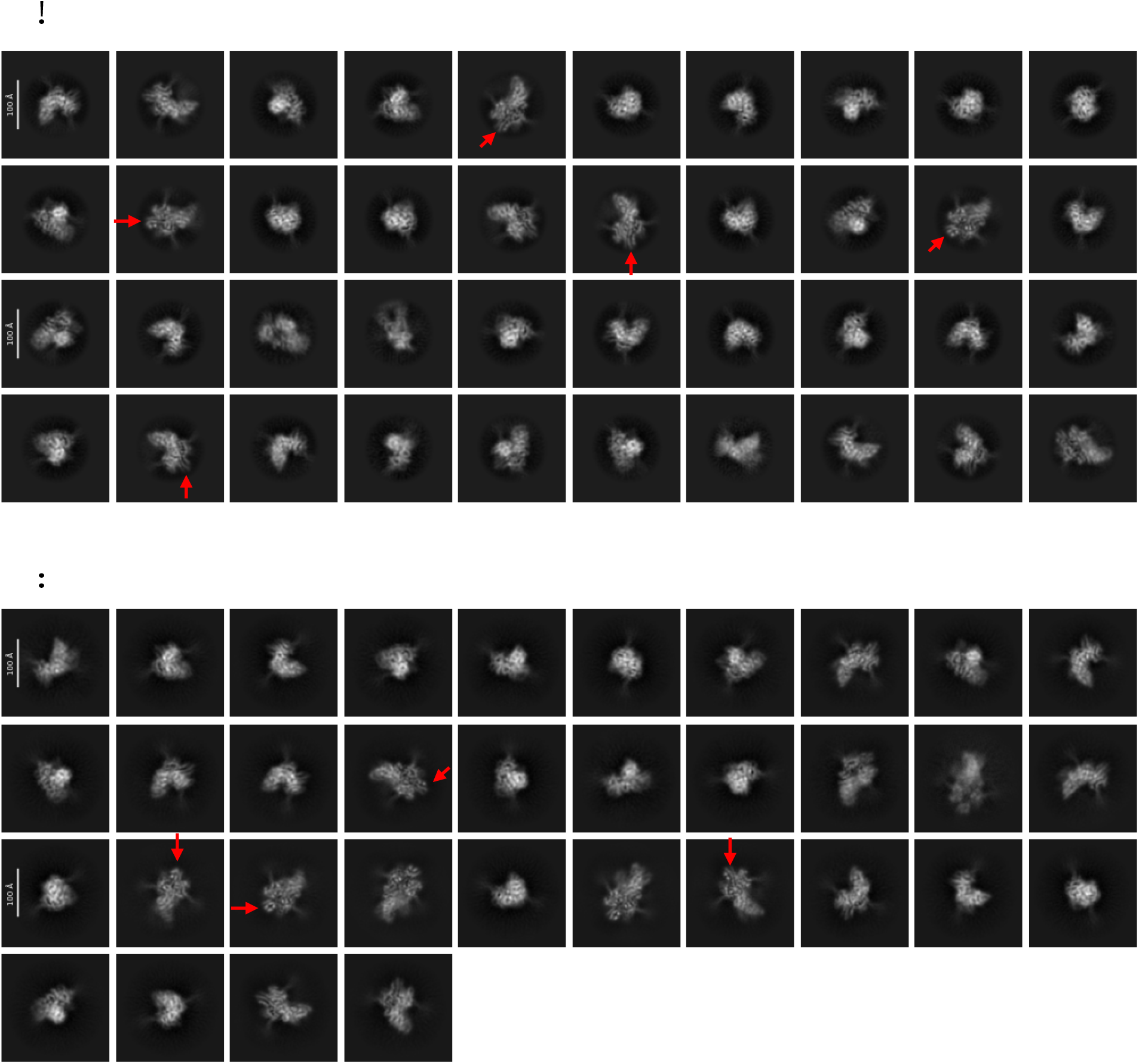
2D class averages for the particles that went into the consensus reconstruction of the HSP90**•**H2 (**A**) and HSP90**•**H4 complex maps (**B**). Arrows point to the binders readily identifiable in a number of 2D classes.

**Table S1.** Data collection, processing and model refinement.

|  |  |  |
| --- | --- | --- |
| **PDB ID/**<br>**EMDB ID** | 13DW / EMD-77013 | 13DX / EMD-77014 |
| **Molecule** | HSP90 with Binder H2 | HSP90 with Binder H4 |
| **Data collection and processing** |  |  |
| Magnification | 105000 |  |
| Voltage (kV) | 300 |  |
| Electron exposure (e-/Å<sup>2</sup>) | 50 |  |
| Defocus range (µm) | -1.0 ~ -3.0 |  |
| Pixel size (Å) | 0.41335 (super-resolution) |  |
| Symmetry imposed | C1 |  |
| Initial particle images (no.) | 1334200 | 263300 |
| Final particle images (no.) | 435700 | 127300 |
| Map resolution (Å) | 2.93 | 3.06 |
| FSC threshold | 0.143 | 0.143 |
| **Refinement** |  |  |
| Initial model used as templet |  | In-silico model |
| **Model composition** |  |  |
| Non-hydrogen atoms | 11360 | 11358 |
| Protein residues | 1390 | 1390 |
| Ligands | 4 | 4 |
| **B factors (Å)**<br>**Min/max/mean** |  |  |
| Protein | 17.29/182.80/79.00 | 21.51/180.52/89.95 |
| Ligand | 16.05/41.26/27.68 | 33.15/56.65/40.16 |
| **R.m.s. deviations** |  |  |
| Bond lengths (Å) | 0.002 | 0.002 |
| Bond angles (°) | 0.486 | 0.506 |
| **Validation** |  |  |
| MolProbity score | 2.42 | 2.36 |
| Clashscore | 12.74 | 13.77 |
| Poor rotamers (%) | 3.75 | 2.71 |
| CaBLAM outliers (%) | 3.87 | 5.01 |
| EMRinger score | 2.00 | 1.39 |
| Favored (%) | 94.57 | 94.44 |
| Allowed (%) | 5.43 | 5.56 |
| Outliers (%) | 0 | 0.00 |

## References

1. Picard, D., Heat-shock protein 90, a chaperone for folding and regulation. Cell Mol Life Sci, 2002. 59(10): p. 1640–8.

2. Schopf, F.H., M.M. Biebl, and J. Buchner, The HSP90 chaperone machinery. Nat Rev Mol Cell Biol, 2017. 18(6): p. 345–360.

3. Taipale, M., D.F. Jarosz, and S. Lindquist, HSP90 at the hub of protein homeostasis: emerging mechanistic insights. Nat Rev Mol Cell Biol, 2010. 11(7): p. 515–28.

4. Biebl, M.M. and J. Buchner, Structure, Function, and Regulation of the Hsp90 Machinery. Cold Spring Harb Perspect Biol, 2019. 11(9).

5. Trepel, J., et al., Targeting the dynamic HSP90 complex in cancer. Nat Rev Cancer, 2010. 10(8): p. 537–49.

6. Chiosis, G., et al., Structural and functional complexity of HSP90 in cellular homeostasis and disease. Nat Rev Mol Cell Biol, 2023. 24(11): p. 797–815.

7. Li, L., et al., Heat Shock Protein 90 Inhibitors: An Update on Achievements, Challenges, and Future Directions. J Med Chem, 2020. 63(5): p. 1798–1822.

8. Goel, B., S. Jaiswal, and N. Tripathi, Recent advances in HSP90 inhibitors as targeted cancer therapy: Chemical scaffolds, isoform selectivity, and clinical progress. Bioorg Chem, 2025. 163: p. 108782.

9. Kumar, A., et al., Recent progress in the development of HSP90 inhibitors: structure-activity relationship and biological evaluation studies. Mol Divers, 2025.

10. Gomez-Pastor, R., E.T. Burchfiel, and D.J. Thiele, Regulation of heat shock transcription factors and their roles in physiology and disease. Nat Rev Mol Cell Biol, 2018. 19(1): p. 4–19.

11. Wang, L., et al., Targeting Chaperone/Co-Chaperone Interactions with Small Molecules: A Novel Approach to Tackle Neurodegenerative Diseases. Cells, 2021. 10(10).

12. Abramson, J., et al., Accurate structure prediction of biomolecular interactions with AlphaFold 3. Nature, 2024. 630(8016): p. 493–500.

13. Pacesa, M., et al., One-shot design of functional protein binders with BindCraft. Nature, 2025. 646(8084): p. 483–492.

14. Baek, M., et al., Accurate prediction of protein structures and interactions using a three-track neural network. Science, 2021. 373(6557): p. 871–876.

15. Humphreys, I.R., et al., Computed structures of core eukaryotic protein complexes. Science, 2021. 374(6573): p. eabm4805.

16. Wohlwend, J., et al., Boltz-1 Democratizing Biomolecular Interaction Modeling. bioRxiv, 2025.

17. Srivastava, D., et al., Unique interface and dynamics of the complex of HSP90 with a specialized cochaperone AIPL1. Structure, 2023. 31(3): p. 309–317 e5.

18. Chaudhury, S., S. Lyskov, and J.J. Gray, PyRosetta: a script-based interface for implementing molecular modeling algorithms using Rosetta. Bioinformatics, 2010. 26(5): p. 689–91.

19. Kortemme, T. and D. Baker, A simple physical model for binding energy hot spots in protein-protein complexes. Proc Natl Acad Sci U S A, 2002. 99(22): p. 14116–21.

20. Lee, K., et al., The structure of an Hsp90-immunophilin complex reveals cochaperone recognition of the client maturation state. Mol Cell, 2021. 81(17): p. 3496–3508 e5.

21. Lavery, L.A., et al., Structural asymmetry in the closed state of mitochondrial Hsp90 (TRAP1) supports a two-step ATP hydrolysis mechanism. Mol Cell, 2014. 53(2): p. 330–43.

22. Bepler, T., et al., Positive-unlabeled convolutional neural networks for particle picking in cryoelectron micrographs. Nat Methods, 2019. 16(11): p. 1153–1160.

23. Gyawali, R., et al., CryoSegNet: accurate cryo-EM protein particle picking by integrating the foundational AI image segmentation model and attention-gated U-Net. Brief Bioinform, 2024. 25(4).

24. Ali, M.M., et al., Crystal structure of an Hsp90-nucleotide-p23/Sba1 closed chaperone complex. Nature, 2006. 440(7087): p. 1013–7.

25. Richter, K., S. Walter, and J. Buchner, The Co-chaperone Sba1 connects the ATPase reaction of Hsp90 to the progression of the chaperone cycle. J Mol Biol, 2004. 342(5): p. 1403–13.

26. McLaughlin, S.H., et al., The co-chaperone p23 arrests the Hsp90 ATPase cycle to trap client proteins. J Mol Biol, 2006. 356(3): p. 746–58.

27. Djuzenova, C.S., et al., Dual PI3K- and mTOR-inhibitor PI-103 can either enhance or reduce the radiosensitizing effect of the Hsp90 inhibitor NVP-AUY922 in tumor cells: The role of drug-irradiation schedule. Oncotarget, 2016. 7(25): p. 38191–38209.

28. Synoradzki, K., et al., Interaction of the middle domains stabilizes Hsp90alpha dimer in a closed conformation with high affinity for p23. Biol Chem, 2018. 399(4): p. 337–345.

29. Simpson, N.E., et al., High levels of Hsp90 cochaperone p23 promote tumor progression and poor prognosis in breast cancer by increasing lymph node metastases and drug resistance. Cancer Res, 2010. 70(21): p. 8446–56.

30. Miyata, Y., H. Nakamoto, and L. Neckers, The therapeutic target Hsp90 and cancer hallmarks. Curr Pharm Des, 2013. 19(3): p. 347–65.

31. Punjani, A., et al., cryoSPARC: algorithms for rapid unsupervised cryo-EM structure determination. Nat Methods, 2017. 14(3): p. 290–296.

